# “Um…, it’s really difficult to… um… speak fluently”: Neural tracking of spontaneous speech

**DOI:** 10.1101/2022.09.20.508802

**Authors:** Galit Agmon, Manuela Jaeger, Reut Tsarfaty, Martin G Bleichner, Elana Zion Golumbic

## Abstract

Spontaneous real-life speech is imperfect in many ways. It contains disfluencies and ill-formed utterances and has a highly variable rate. When listening to spontaneous speech, the brain needs to contend with these features in order to extract the speaker’s meaning. Here, we studied how the neural response is affected by four specific factors that are prevalent in spontaneous colloquial speech: (1) the presence of non-lexical fillers, (2) the need to detect syntactic boundaries in disfluent speech, and (3) variability in speech rate. Neural activity (EEG) was recorded from individuals as they listened to an unscripted, spontaneous narrative, which was analyzed in a time-resolved fashion to identify fillers and detect syntactic boundaries. When considering these factors in a speech-tracking analysis, which estimates a temporal response function (TRF) to describe the relationship between the stimulus and the neural response it generates, we found that the TRF was affected by both of them. This response was observed for lexical words but not for fillers, and it had an earlier onset for opening words vs. closing words of a clause and for clauses with slower speech rates. These findings broaden ongoing efforts to understand neural processing of speech under increasingly realistic conditions. They highlight the importance of considering the imperfect nature of real-life spoken language, linking past research on linguistically well-formed and meticulously controlled speech to the type of speech that the brain actually deals with on a daily basis.

## Introduction

Neural speech tracking has become an increasingly useful tool for studying how the brain encodes and processes continuous speech (Brodbeck & Simon, 2020; Obleser & Kayser, 2019). Importantly, characterizing linguistic attributes of speech on a continuous basis gives a new angle to auditory attention and neurolinguistics, as researchers have been able to dissociate neural responses driven by the acoustics of speech from those capturing higher-order processes in a dynamically changing speech signal, such as phonological identity and semantic expectations (Brodbeck et al., 2018; Gillis et al., 2021; Inbar et al., 2020; Keitel et al., 2018). And yet, the speech stimuli used in these studies are generally highly scripted and edited, taken – for example – from audiobooks or TED talks, which are also extensively rehearsed and are delivered by professionals. These stimuli are in many ways different from colloquial speech that we hear every day (Blaauw, 1994; Face, 2003; Goldman-Eisler, 1968, 1972; Haselow, 2017; Huber, 2007; Mehta & Cutler, 1988). Spontaneous speech is also markedly less fluent than scripted speech, peppered with fillers and pauses, and often includes partial or grammatically incorrect syntactic structures (Auer, 2009; Linell, 1982, Chapter 6)(Haselow, 2017, Chapter 1; Linell, 1982; Shriberg, 2001). Spontaneous speech is also more variable than scripted speech in terms of its speech rate (Goldman-Eisler, 1961; J. L. Miller et al., 1984). Silent pauses in spontaneous speech occur less consistently on syntactic boundaries compared to reading (Goldman-Eisler, 1972; Wang et al., 2010). These differences may render the syntactic analysis of spoken speech less trivial compared to the planned, well-structured sentences in scripted speech materials. By focusing mostly on scripted speech materials, past research might have overlooked important processes which are essential for understanding naturalistic speech.

Addressing this gap, in the current EEG study we assess neural speech tracking of spontaneously generated speech and focus on the specific challenges the brain has to cope with when processing spontaneous speech: the abundant presence of fillers, online segmentation and detection of syntactic boundaries and variability in speech rate.

### Disfluency and fillers

A prominent feature of spontaneous speech is that it contains frequent pauses, self-corrections and repetitions (Bortfeld et al., 2001; Clark & Wasow, 1998; Fox Tree, 1995). These disfluencies are generally accompanied by fillers, which are non-lexical utterances (or filled-pauses) such as “um” or “uh”, or discourse markers such as “you know” and “I mean” (Fox Tree & Schrock, 2002; Tottie, 2014). Fillers can take on different forms, however, their prevalence across a multitude of spoken languages (Tian et al., 2017; e.g. Wieling et al., 2016) suggests it is a core feature of spontaneous speech. Although fillers do not, in and of themselves, contribute specific lexical information, they are also not mere glitches in speech production. Rather, they likely serve several important communicative goals, helping the speaker in transforming their internal thoughts to speech and helping the listener interpret this speech. Specific roles that have been attributed to fillers include signaling hesitation in speech planning (Corley & Stewart, 2008), conveying lack of certainty in the content (S. E. Brennan & Williams, 1995; Smith & Clark, 1993), serving as a cue to focus attention on upcoming words or complex syntactic phrases (Clark & Fox Tree, 2002; Fox Tree, 2001; Fraundorf & Watson, 2011; Watanabe et al., 2008), signaling unexpected information to come (Arnold et al., 2004; Barr & Seyfeddinipur, 2010; Corley et al., 2007) and disambiguating syntactic structures (Bailey & Ferreira, 2003). Moreover, studies have shown that the presence of fillers improves accuracy and memory of speech content (S. E. Brennan & Schober, 2001; Corley et al., 2007; Fox Tree, 2001; Fraundorf & Watson, 2011) and that surprise-related neural responses (event-related potentials; ERPs) to target words are reduced if they are preceded by a filler (Collard et al., 2008; Corley et al., 2007). And yet, despite their clearly important role in the production and perception of spontaneous speech, fillers and other disfluencies are generally absent in planned speech that is used in most lab speech-tracking experiments. Therefore, how the brain processes fillers has not been studied extensively.

### The unorganized nature of spontaneous sentences

Unlike written text or highly edited spoken scripts, spontaneous speech is constructed ‘on the fly’, and represents the speakers unedited and somewhat ‘unpolished’ internal train of thought. As a consequence, spontaneous speech does not always contain clear sentence endings, is not always grammatically correct, and sentences can seem extremely long and less concise (e.g. *“and, so, then we went to the bus, but it, like, didn’t come, the bus back home I mean, so we had to wait, I really don’t know for how long”*) (Auer, 2009; Halliday, 1989, Chapter 6; Haselow, 2017; Linell, 1982). This poses a challenge to the listener of how to parse the continuous input-stream into meaningful syntactic units correctly.

Syntactic parsing is the process of analyzing the string of words into a coherent structure, and is critical for speech comprehension, even under assumptions of a “good-enough” or noisy parse (Ferreira & Patson, 2007; Traxler, 2014). To detect syntactic boundaries in spoken language, listeners rely on their accumulated syntactic analysis of an utterance as well as on prosodic cues such as pauses and changes in pitch and duration (Ding et al., 2016; Fodor & Bever, 1965; Garrett et al., 1966; Har-shai Yahav & Zion Golumbic, 2021; Hawthorne & Gerken, 2014; Kaufeld et al., 2020; Langus et al., 2012; Strangert, 1992; Strangert & Strangert, 1993). There are some behavioral and neural indications that words occurring at the final position of syntactic structures have a ‘special status’. In production, words in final positions tend to be prosodically marked (Cooper & Paccia-Cooper, 1980; Klatt, 1975). In comprehension, reading studies show that sentence-final words have prolonged reading and fixation times as well as increased ERP responses. These effects are known as ‘wrap up effects’, which is a general name for the integrative processes triggered by the final word (Just and Carpenter, 1980; Stowe et al., 2018 for review). Independently, there is also evidence for neural responses associated with detecting prosodic breaks (Pannekamp et al., 2005; Peter et al., 2014; Steinhauer et al., 1999), which often coincide with syntactic boundaries. However, the neural correlates of syntactic boundaries have seldom been studied in the context of spoken language, and particularly not for spontaneous speech, where sentence boundaries are not as well-formed as in scripted language materials.

### Speech Rate

Another characteristic of spontaneous speech studied here is speech rate across different sentences. Spontaneous speech is produced ‘on-the-fly’, which can yield speech that at times is highly coherent, fast and excited, and at times is prolonged and interspersed with pauses and hesitations (Goldman-Eisler, 1961, 1972; J. L. Miller et al., 1984). Generally, higher speech-rate mean that information needs to be integrated in a shorter amount of time, which imposes higher cognitive load on the listener and can affect the processing of speech in many ways. For example, processing compressed speech decreases speech comprehension and intelligibility (Ahissar et al., 2001; Ahissar & Ahissar, 2005; Chan & Lee, 2005; Vaughan & Letowski, 1997; Verschueren et al., 2022). Additionally, phonological and lexical decoding are affected by speech rate and dynamically adjusted to local changes in speech rate (Dilley & Pitt, 2010; Dupoux & Green, 1997; Joanne L. Miller et al., 1986). Several studies have shown that neural tracking of continuous speech can be affected by artificially manipulating speech-rate (Ahissar et al., 2001; J. A. Müller et al., 2019; Verschueren et al., 2022). However, few have looked at the natural variations in speech rate in spontaneous discourse.

To summarize, spontaneous speech differs in many ways from scripted speech. Here we focused on three key characteristics of spontaneous speech and ask how the presence of fillers, the need to detect syntactic boundaries on line and the natural variations in speech-rate affect listeners’ neural response to speech. To do so, we analyzed EEG-recorded neural responses from individuals listening to a six-minute long monologue rich in those features, recorded from a speaker spontaneously recounting a personal experience. We analyzed the monologue to identify fillers and syntactic boundaries and estimate speech-rate, which were used as data-driven regressors for analyzing the neural activity, using multivariate speech-tracking analysis of the EEG data. In doing so, this study aims to bridge the gap between the vast literature studying brain responses to meticulously controlled speech and language materials, and the type of speech materials that the brain deals with on a regular daily basis.

## Methods

### Participants

Twenty participants took part in this experiment. All participants were native speakers, right-handed (16:4 F:M; age range 20-30, average 23.1±2.65). Prior to their participation, participants signed informed consent approved by the IRB of the University [anonymized as per journal policy] and were compensated for their participation with credit or payment.

### Stimuli and procedure

The stimulus was a single 6-minute long recording of a personal narrative, told in the first person by a female native speaker. The narrative was neutral, unscripted, and spontaneously generated, and described the speaker’s participation in a Facebook group called “stories with no point” and a social face-to-face meet-up organized by members of the group. The only instructions given to participants were to listen passively to the story presented without breaks. This 6-minute session was used as an interlude between two parts of another experiment (currently unpublished) that focused on changes in low-level auditory responses over time and is orthogonal in its goals to the data reported here.

### Linguistic analysis of the speech stimulus

The speech narrative was transcribed manually by two independent annotators who were native speakers of the language (anonymized, as per the journal’s double-blind peer review policy), and verified by an expert linguist. The onset and duration of each word and of fillers were identified and time-stamped by two independent annotators, and confirmed by a third using the software Praat (Boersma & Weenink, 2021).

Based on the transcription, an expert linguist parsed the speech stimulus into major syntactic units, most notably clauses. We marked boundaries of all main clauses, defined as the minimal unit containing a predicate and all its complements and modifiers. This includes clauses in coordinate constructions (starting with “and” or “but”). In many cases, a clause can contain a subordinate clause, whose boundaries we also marked. These included complement clauses (e.g., “[we decided [*that we would meet in one of the gardens*]]”), adverbial clauses (e.g., “[we met [*because we needed to talk*]]”) and relative clauses (e.g., “[we met a delegation [*that came from Japan*]]”). We also marked the boundaries of heavy phrases such as appositives (e.g., “[We decided to go there, [*me and my friends*]]”) or ellipsis (e.g., “[I was fifteen years old then, [*in the tenth grade*]]”).

Although syntactic parsing was done primarily based on the speech transcript, it was double-checked relative to the audio-recording in search of cases where the spoken prosody suggested a different intended parsing than the textual syntactic analysis. For example, the text “I went home with my friends” could be considered a single clause from a purely textual perspective. However, in the audio recording, the speaker inserted a pause after the word “home”, and hence, a listener would likely have identified the word “home” as the final word in the clause, before the speaker decided to continue it with a prepositional phrase (“with my friends”). Due to this perceptual consideration, in such cases, we marked both the word “home” and the word “friends” as the final word in the clause.

The time-stamped transcription and syntactic parsing were used to annotate the speech stimulus according to the following four word-level features s, which were used to analyze the neural response (described below and summarized in Figure 1):

1. *Fillers vs. non-fillers:* Fillers were identified and time-stamped as part of the transcription process. The definition of fillers included both filled-pauses (e.g., “um”, “uh”); and filler discourse markers, which are words that are lexical units, but their use in the utterance is not related to their original lexical meaning; (e.g., “like”, “well”). A total of 92 fillers were detected in the speech stimulus.
2. *Words at syntactic boundaries (opening vs. closing words)*: Opening and closing words were identified based on the marking of syntactic clause boundaries described above. There were a total of 166 opening words and 142 closing words (since some clauses were syntactically incomplete and did not have a clear closing word).
3. *Word Length (short vs. long words):* The length of each word was evaluated based on the time-stamped transcription and the median length was used to distinguish between short and long words (median length: 319 ms).
4. *Information Content (function vs. content words):* We also differentiated between words that carry the most information (content words; defined as nouns, verbs, adjectives and adverbs) vs. words that mostly play a syntactic role (function words; defined as pronouns, auxiliary verbs, prepositions, conjunctions and determiners). In this data set, there were 413 content and 250 function words.

**Figure 1.**
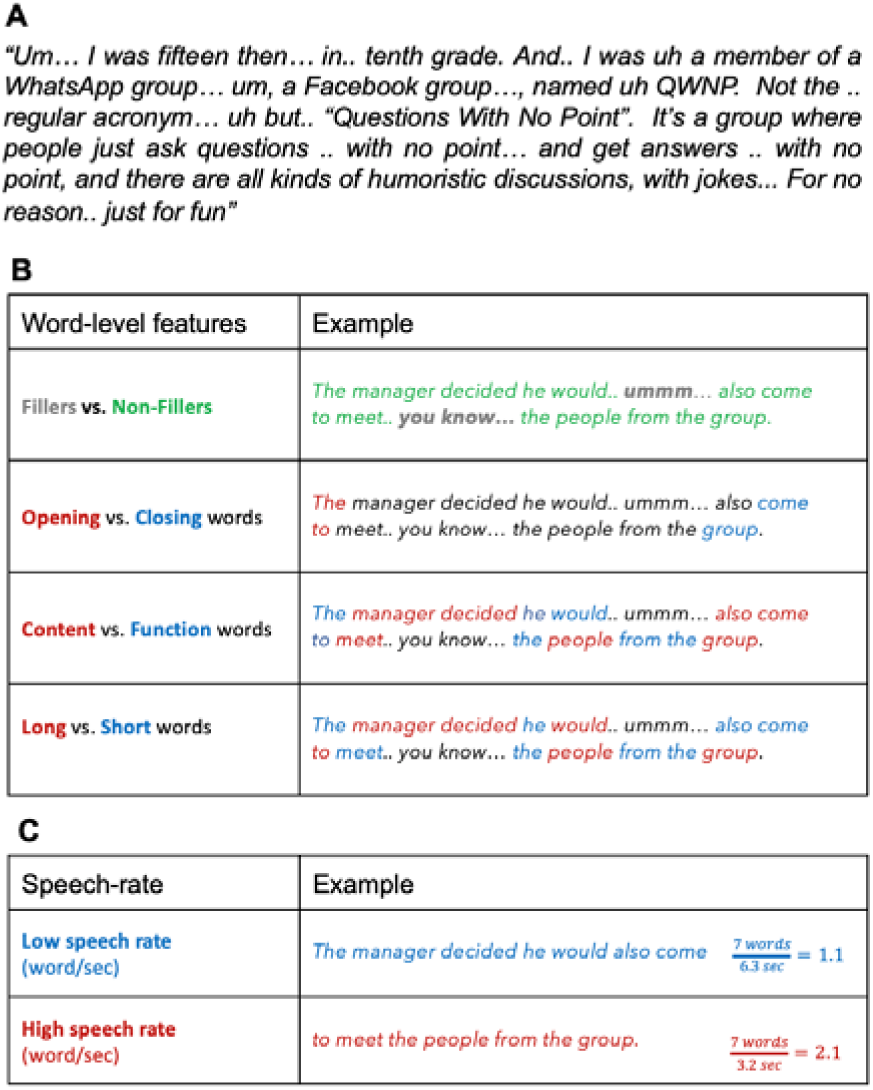
**A.** Excerpt from the speech stimulus used, demonstrating the disfluencies of spontaneous speech. **B.** Example of the four word-level features of spontaneous speech analyzed here. **C.** Example of the quantification of speech-rate at the clause level.

**Figure 2.**
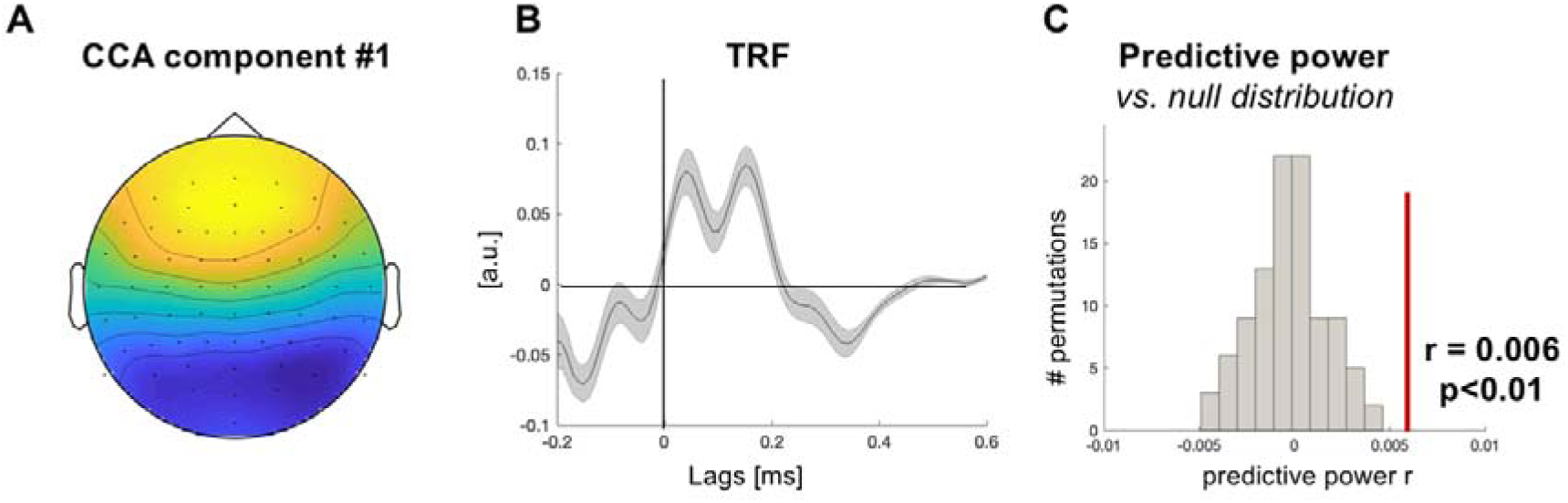
**A.** The scalp distribution of CCA component #1, which corresponds to the expected auditory scalp topography. **B.** The TRF estimated for CCA component #1. **C.** The predictive power of the TRF shown in B was significantly larger than could be obtained by chance, shown here relative to the null-distribution of predictive power from 100 randomly shuffled S-R combinations.

### Analysis of Speech-Rate

Potential effects of variability in speech-rate cannot be assessed at a single-word level, but require assessing the rate of speech over longer periods of times. Here we chose to quantify the mean speech-rate within each main clause (including any embedded complement clauses and restrictive relative clauses), following the rationale that a clause is the basic unit over which information needs to be integrated during on-line listening.

We tested two metrics for operationalizing speech-rate: syllable-rate and word-rate. These metrics are highly correlated with each other (r = 0.76, p<10^−11^ in the current data set), but emphasize slightly different aspects of information-transfer, with syllable-rate capturing the rate of acoustic input and word-rate capturing the rate of linguistic input and general fluency. The word-rate of each clause was quantified as the number of words in a clause (not including fillers) divided by its length. Similarly, the syllable-rate quantified as the number of syllables in a clause (not including fillers) divided by its length.

### EEG recordings

EEG was recorded using a 64 Active-Two system (BioSemi) with Ag-AgCl electrodes, placed according to the 10-20 system, at a sampling rate of 1024 Hz. Additional external electrodes were used to record from the mastoids bilaterally and both vertical and horizontal EOG electrodes were used to monitor eye movements. The experiment was conducted in a dimly lit, acoustically, and electrically shielded booth. Participants were seated on a comfortable chair and were instructed to keep as still as possible and breathe and blink naturally. Experiments were programmed and presented to participants using PsychoPy (https://www.psychopy.org) (Peirce et al., 2019).

### EEG preprocessing and speech-tracking analysis

EEG preprocessing and analysis were performed using the matlab-based FieldTrip toolbox (Oostenveld et al., 2011) as well as custom-written scripts. Raw data was first visually inspected, and time-points with gross artifacts exceeding ±50 μV (and were not eye movements) were removed. Independent Component Analysis (ICA) was performed to identify and remove components associated with horizontal or vertical eye movements as well as heartbeats (Onton et al., 2006). Any remaining noisy electrodes that exhibited either extreme high-frequency activity (>40Hz) or low-frequency activity/drifts (<1 Hz), were replaced with the weighted average of their neighbors using an interpolation procedure. The clean EEG data was filtered between 1-10Hz. The broadband envelope of the speech was extracted using an equally spaced filterbank between 100 to 10000 Hz based on Liberman’s cochlear frequency map (Liberman, 1982). The narrowband filtered signals were summed across bands after taking the absolute value of the Hilbert-transform for each one, resulting in a broadband envelope signal. The speech-envelope and EEG data were aligned in time and downsampled to 100Hz, for computational efficiency.

To increase the signal-to-noise ratio of the EEG data and reduce the dimensionality of the data, we applied Correlated Component Analysis (CCA) to the clean EEG data. This procedure identifies spatio-temporal components with high inter-subject correlation, and is particularly effective for experiments studying neural responses to continuous natural stimuli (Parra et al. 2018, Dmochowski et al. 2015; https://github.com/dmochow/rca). We performed speech-tracking analysis on the first three CCA components, but ultimately only the first CCA component showed above-chance speech-tracking responses (see below), therefore subsequent analyses were limited only to this component. In the supplementary material (S1) we report results of identical speech-tracking analysis performed on the original EEG data from all 64-channels, to demonstrate that these are qualitatively similar to those obtained using the CCA component, but statistical evaluation of effects was restricted only to the CCA data, to avoid unnecessary multiple comparisons.

Speech-tracking analysis was performed using scripts from the STRFpak Matlab toolbox (strfpak.berkeley.edu) which were adapted for EEG data (Zion Golumbic, Cogan, et al. 2013, Har-shai Yahav 2021). In this approach, a normalized reverse correlation is applied in order to estimate a Temporal Response Functions (TRF) that best describes the linear relationship between features of the presented speech stimulus S(t) and the neural response R(t). Speech tracking analysis was performed separately for each of feature of interest, as described below.

### (1) Acoustic-only Model

The acoustic-only model was estimated to describe the linear relationship between the broadband envelope of the speech stimulus (S) and the neural response (R), derived from the EEG data (R). For S we used the broadband envelope of the speech, which was estimated by first bandpass filtering the speech stimulus into 10 equally spaced bands between 100 to 10000 Hz based on Liberman’s cochlear frequency map, taking the amplitude of each narrow-band using the absolute value of the Hilbert transform, and then summing across bands. For R we used the CCA component (1-20Hz). This analysis was also repeated for filtered responses in the canonical delta band (0.5-3Hz) and theta band (4-8Hz; zero-phase FIR filter), and are reported as supplementary data (Figure S2). S and R were segmented into 12 equal-length non-overlapping epochs (~30 seconds each). To minimize the effects of over-fitting, TRFs were estimated using a jackknife leave-one-out cross-validation procedure, whereby TRFs are estimated using a 5-fold train-test procedure. In each iteration, the model was trained on 10 epochs and tested on the remaining 2 epochs that were not included in the training. To select the optimal regularization tolerance value, in each fold of the training, we performed a jackknife leave-one-out cross-validation procedure in which TRFs were estimated for a range of regularization values [0.005-0.01] using all-but-one of the training trials (i.e. 9 trials), and the Pearson’s correlation (r) was calculated between the estimated and actual EEG signal for the 1 left-out validation trial. After this procedure was repeated for all possible jackknifes (10 times), the tolerance level that yielding the highest r-value across all jackknifes was selected, and the estimated TRFs for that tolerance level were averaged across jackknifes. The predictive power of the resulting TRF was then assessed using the test data (the 2 trials that were not included in the training), by comparing the EEG signals of the test-data with the predicted response using the TRF estimated from the training data, and calculating the Pearson’s correlation (r) between them. This entire procedure was repeated 5 times, using different partitioning of the data into independent training-testing data sets, and the mean predictive power and TRFs across these 5 folds are reported. The optimized tolerance values estimated for the acoustic-only model were then used in all subsequent analyses.

Statistical significance of the speech tracking response using the acoustic-only model was evaluated using a permutation test as follows: We repeated the TRF estimation procedure on 100 different permutations of mismatched S-R pairs (S from one segment with R from another). This yielded a null-distribution of predictive power values that could be obtained by chance. The predictive power of the correct S-R pairing was compared to this distribution and was considered significant if it fell in the top 5% of the null-distribution (one-sided).

### (2) Word-level Models

For the word-level models, the full acoustic envelope was separate into independent regressors containing only the envelope of words/utterances corresponding to a particular feature. We then performed a series of multivariate TRF analyses to test different word-level research questions, as follows:

#### Fillers vs. non-filler model

Comparing a regressor representing all the fillers vs. a regressor with non-filler words (Figure 3A). Since non-filler words were extremely more prevalent than filler words, a subset of non-filler words were randomly selected for this analysis to match the number of fillers. To avoid selection-bias, this analysis was conducted 10 times using different sub-selections of non-filler words.

**Figure 3.**
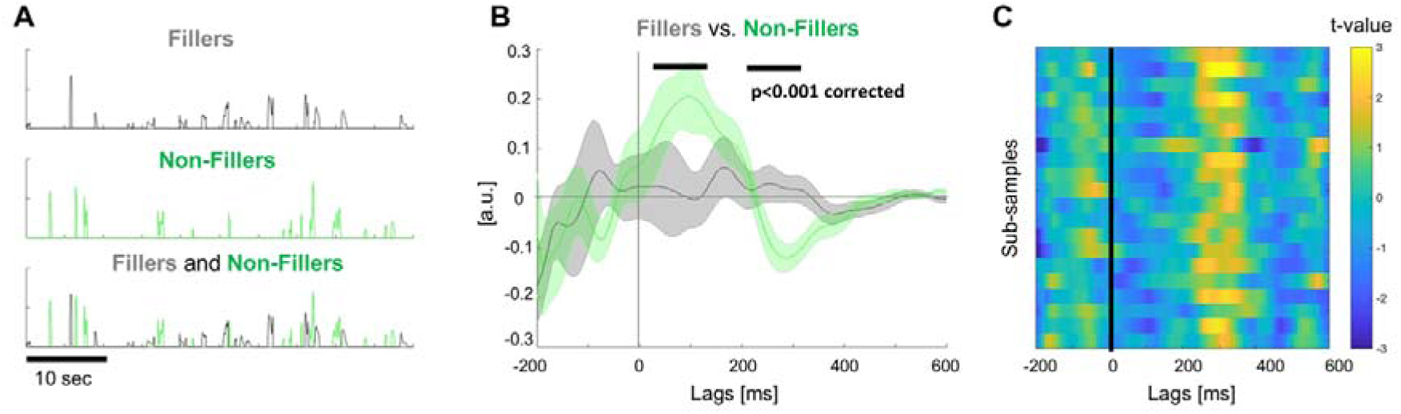
**A.** Illustration of the regressors use to assess TRFs for fillers vs. non-filler words. Top two rows: Each regressor contains only the envelope of the relevant portion of the stimulus. Bottom row: the two regressors overlaid. For the non-filler regressor, 92 words were randomly sampled, to equate with the number of fillers. **B.** TRFs for fillers vs. non-fillers, which differed significantly in both an early (20-120ms) and late time window (220-360ms; p<0.001). **C.** Results from 20 different repetitions of this analysis using different sub-samples of non-filler words. Figure shows the t-value at each time-lag reflecting the difference between TRFs for the filler vs. the non-filler regressors.

#### Syntactic Boundary model

Comparing a regressor with opening vs. closing words (Figure 4A). Note that although the research question addressed in this analysis was the difference between opening vs. closing words, mid-sentences words were also included in this model as a separate regressor, so as to maintain a full representation of the entire speech envelope for the purpose of cross-validation. However, the TRF for mid-sentence words was not compared statistically to the others, since they were found to be too variable acoustically.

**Figure 4.**
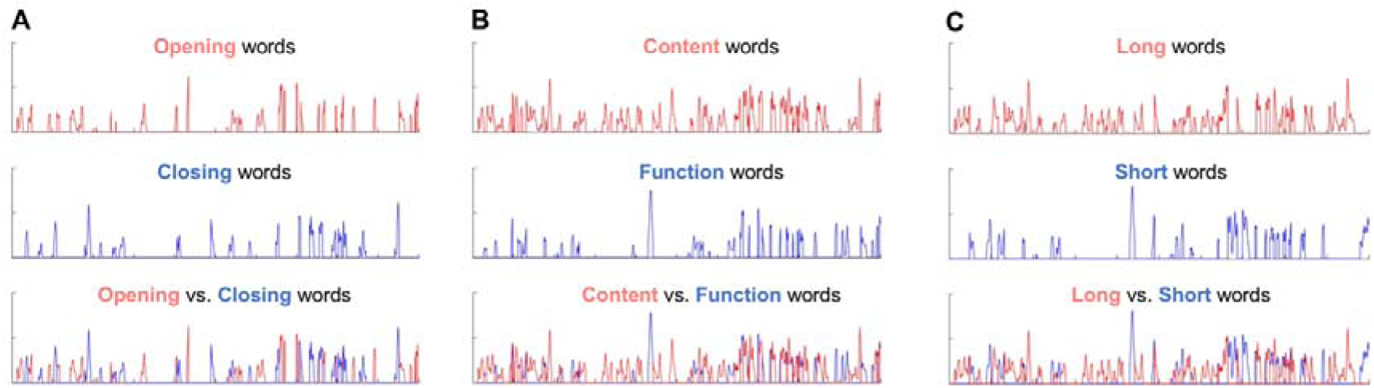
Illustration of the regressors use to assess TRFs for **A.** opening vs. closing words. **B.** content vs. function words. **C.** long vs. short words (median split of stimulus). The top two rows in each panel show the individual regressors, that contains only the envelope of the relevant portion of the stimulus. Bottom row: the two regressors overlaid. For B&C this constitutes the full stimulus.

We also tested two control models, to test potential alternative explanation for differences between fillers/non-fillers or opening/closing words. *Information Content model:* Comparing a regressor with content vs. function words (Figure 4B). *Word length model:* Comparing a regressor with short vs. long words (median split; Figure 4C). The fillers-regressor was included in all word-level models, to allow us to compare TRFs to fillers vs. words with specific features.

Each model was tested using a multivariate TRF analysis, that estimates a separate TRF for each regressor, using the 5-fold train-test regime described above. We then compared the amplitude of the TRFs estimated for the different regressors in each model, using a pair t-test at each time point, and corrected for multiple corrections over time using a global-statistic permutation test (1,000 permutations).

### (3) Speech-rate Models

Speech rate was estimated for each clause (as described above) and clauses were divided into two groups corresponding to “low”/“high” speech rate. This was initially done using a median split of speech rates, but given that clauses with faster rates may be shorter than clauses with lower speech rates, the cutoff for separating the two was shifted slightly, to ensure that each group represented roughly half of the full stimulus (cutoff values: syllable rate: 5.23 syllable/sec; word rate 2.14 words/sec; solid line in Figure 6A). Using this criterion, clauses with low speech rate had a mean syllable rate of 4.16±0.67 syllable/sec and a mean word rate of 1.72±0.25 words/sec; and clauses with high speech rate had a mean syllable rate of 7.36±1.42 syllable/sec and a mean word rate of 3.23±0.81 words/sec. Two separate regressors were created, representing the envelope of the clauses with low vs. high speech rate (Figure 6B). A multivariate TRF analyses was performed using the two regressors for each feature, using the 5-fold train-test regime described above. We then tested for significant differences between the TRFs estimated for low vs. high values of each feature, using a pair t-test at each time point, and corrected for multiple corrections over time using a global-statistic permutation test (1,000 permutations).

**Figure 5.**
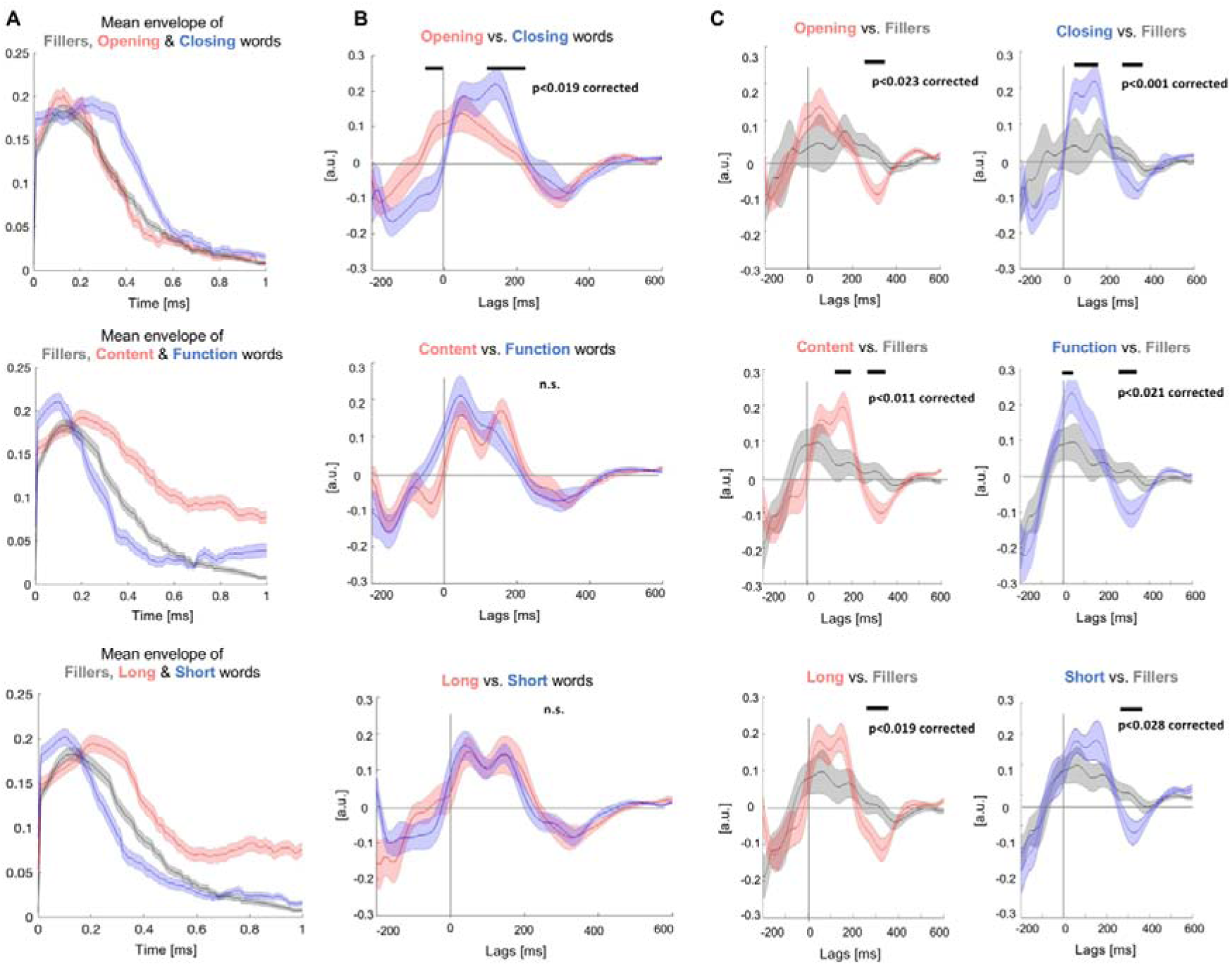
**A.** The average envelope across words\fillers, for the features tested in each word-level model. This comparison indicated that the mean amplitude (volume) was relatively similar across word-level features, however closing words, content words and long words were more prolonged relative to opening words, function words and short words, respectively. **B.** The TRFs estimated in each word-level model. These results revealed significant differences in the neural response to opening vs. closing words (p<0.019, corrected), but not differences between content vs. function words or between long vs. short words. **C.** Comparison of the TRFs to different types of words vs. fillers (estimated as part of each multivariate word-level model). In all cases, significant differences were observed for the late negative TRF response (~250-400ms), and sometimes for the early positive TRF responses as well (~50-180ms).

**Figure 6.**
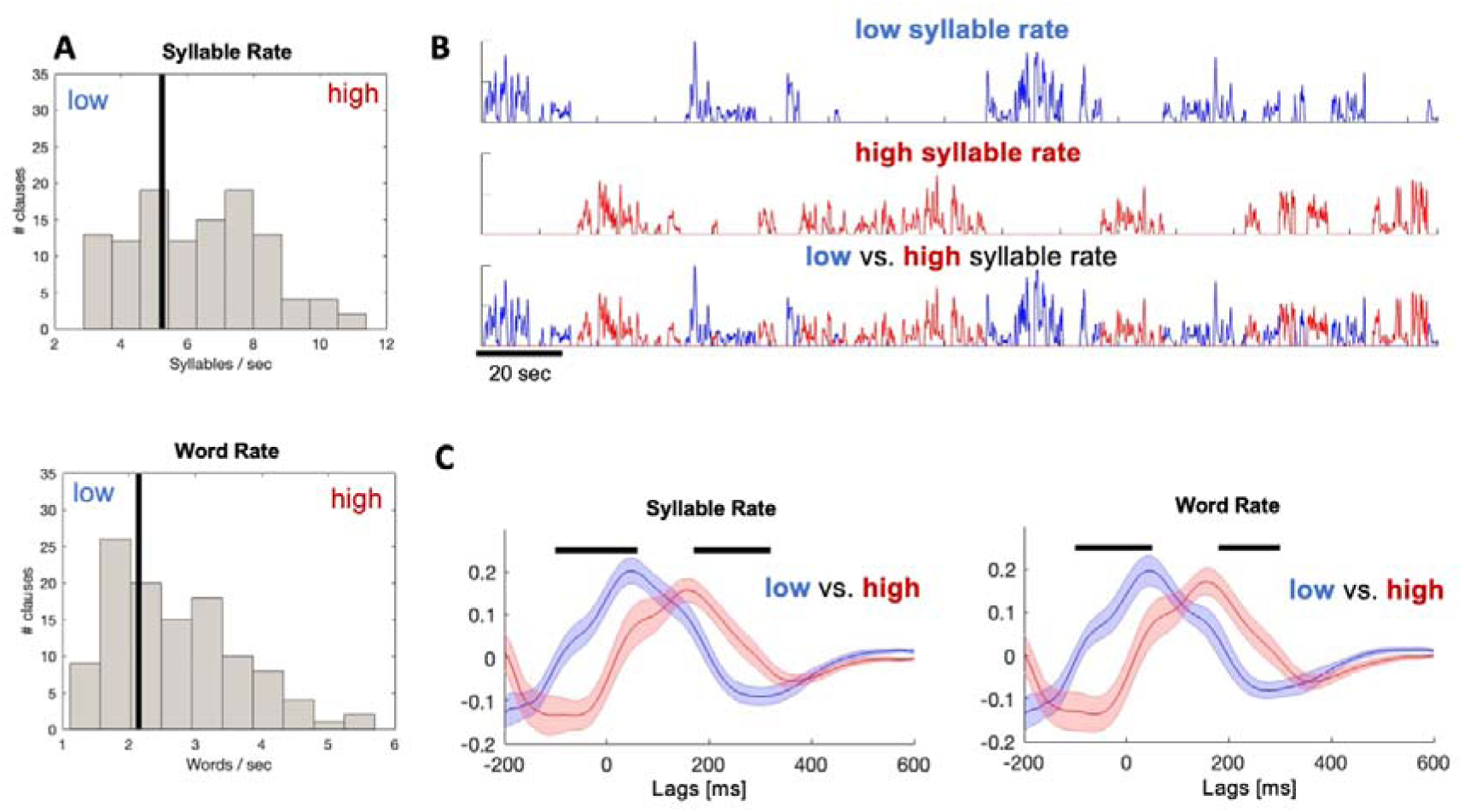
**A.** The distribution of syllabe-rate (top) and word-rate values (bottom) across all clauses. The vertical line in each panel indicates the cutoff value used to split the clauses into low/high speech rate (median split of the stimulus). **B.** Excerpts demonstrating the time-course of the regressors representing clauses with low/high syllable rate. The top two rows in each panel show the individual regressors, and the bottom row shows both regressors overlaid, which together constitute the full stimulus. **C.** The TRFs estimated for clauses with low (blue) vs. high (red) speech rate, shown separately for the two operationalization of this construct (syllable rate and word rate). Horizonal black lines indicate the time windows where significant differences were observed between the TRFs (p<0.001, corrected).

## Results

### Acoustic-only model

The topography of the first CCA component corresponded to the traditional EEG topography for auditory responses (Figure 2 and Figure S1), and this was the only CCA component that showed a significant speech-tracking response (r = 0.006; p<0.01 relative to permutations; Figure 2C). Therefore, all subsequent analyses were restricted only to this component. The acoustic TRF showed a positive deflection between 0-200ms (with several local peaks riding on), and a negative deflection between 300-400ms. This temporal profile is in line with TRF typically obtained for speech stimuli, although the precise timing and modulation depth of different peaks can be highly dependent on the specific regularization parameters selected and the characteristics of the regressor used.

### Word-level models

When comparing the TRF for fillers vs. non-filler words (Figure 3) we found that the TRFs for fillers was extremely flat and was significantly lower than TRF for non-filler words in the two time-windows surrounding peaks in the speech-tracking response (20-120ms and between 220-360ms; p<0.001 corrected). This analysis was repeated for 20 different sub-samples of non-lexical words (proportion matched to the fillers) which yielded similar results, indicating the robustness of this results (Figure 3C and see also Figure 5C).

The other word-level regressors tested included comparison of the TRFs derived for opening vs. closing words, content vs. function words and long vs. short words (Figure 4 & 5), and the TRFs to each word-type were also compared to TRFs for fillers.

Alongside the TRF analysis comparing the neural responses to different word-level features, we also evaluated the acoustic differences between the different type of words, by calculating the average envelope amplitude across words and calculating their duration, for each feature (Figure 5A). This is important in order to determine whether differences in neural responses might be due to systematic differences in the acoustic properties of different word-types. This revealed that closing words had a similar mean amplitude relative to opening words, but were more prolonged (median duration 485±164 ms vs. 300±208 ms, p<10^10^). This lengthening for closing words has been previously documented in final positions of major syntactic constituents and shown to be an important cue for segmentation (Klatt, 1975; Langus et al., 2012; Vaissière, 1983). As expected, we also found that content words were more prolonged relative to function words (median duration 400±188 ms vs. 207±136 ms, p<10^10^), however these did not differ significantly in their mean peak amplitude, as was also the case for our separation between long words (median duration 460±160 ms) vs. short words (208±68 ms). Fillers had a similar overall amplitude as all word types, and their duration were comparable to those of the longer words (median duration 481±310 ms).

Interestingly, these differences in acoustics were not reflected in the comparison between the TRFs estimated for each word-level feature. The only significant difference observed was between opening vs. closing words, with the response to closing words peaking later and with a higher amplitude, relative to opening words (Figure 5B top; p<0.019, corrected). However, no differences were found between the neural response to content vs. function words (Figure 5B middle) or between long vs. short words (Figure 5B bottom), suggesting that the word-duration alone probably did not drive the difference between opening vs. closing words. In line with the results showing reduced TRFs for fillers vs. non-filler words in for both the early and late peaks of the TRF (Figure 3), we find that this effect is replicated in all comparisons of TRFs to specific types of words (opening words, closing words, short words, long words, content words and function words) vs. fillers, further supporting the robustness of this result.

### Speech-rate model

As expected, results are highly similar for the two operationalizations of speech-rate: syllable rate and word rate, as they are highly correlated with each other. For both measures we found significant differences between the TRFs for clauses with fast vs. slow speech, which manifested primarily as a latency shift, with earlier response for clauses with slow vs. fast speech. The differences between TRFs were significant in an early time window starting even prior to time-lag 0 (−90-50ms) and in a later time-window (170-300ms; p<0.001 corrected for both, reported time-windows are the overlap between the two measures).

## Discussion

In the current study, we looked at how the neural speech tracking response (TRF) is affected by different features of spontaneous speech. We tested effects of a word’s lexicality (whether it was a filler or not), its role in the structure (whether it opened or closed a syntactic clause) and in the message (whether it was a function or content word), as well as potential differences due to word duration and speech rate. We found that, indeed, the estimated TRFs were modulated by many of these features, suggesting that this response does not merely follow the acoustic of the stimulus but adjusts flexibly to accommodate the linguistic complexity of spontaneous speech. Specifically, we found that fillers (e.g., “um”, “uh”, “like”) elicited a weaker speech-tracking response than words that were part of the core/semantic meaning of the utterance. We also found differences in the TRF latencies for opening vs. closing words in a clause, possibly providing a neural marker for on-line syntactic segmentation. TRFs estimated for entire clauses were also affected by the rate of speech, with earlier latencies found for clauses with slower information-rate. While many findings are in-line with previous research, some of the results are more difficult to interpret mechanistically at this point and require additional follow-up research. However, taken together, these findings highlight the importance of considering the complexities and imperfections of speech when building theoretical models of neural processing of speech, particularly if we strive to understand how the brain contents with the type of speech encountered in actual real life.

### The speech tracking response (TRF)

Before discussing the specific effects found here, it is worth considering what the TRF represents and how it relates to more traditional ERP measures capturing neural responses to language material. The ERP world has a long tradition of associating different deflections in the signal (“components”) to specific perceptual and/or cognitive operations, assigning meaning to both the amplitude and latency of these time-locked responses (Hagoort & Brown, 2000). The recent introduction of speech tracking methods and estimation of TRFs is often thought of as an equivalent to ERPs for continuous stimuli (Brodbeck & Simon, 2020; Dikker et al., 2020). Supporting this notion, TRFs to speech share many commonalities with auditory ERPs, which display similar time-courses of peak and troughs, and similar scalp-topographies and source estimations (Lalor et al., 2009). Early TRF components between 50-150 ms have been shown to be localized to auditory cortex and are primarily driven by the acoustics of the speech (Di Liberto et al., 2015; Ding et al., 2014), whereas linguistic and semantic features such of as predictability, surprise, lexical frequency and semantic, have been shown to affect TRF responses in later time windows between 200-400ms (Brodbeck et al., 2018, 2022; Broderick et al., 2018, 2019; Donhauser & Baillet, 2020; Gillis et al., 2021), in ways reminiscent of the P2 and N400 ERP responses to written words (Kutas & Federmeier, 2011). Hence, it is appealing to interpret the observed TRP responses as an extension of the vast ERP literature to the domain of continuous speech. (Gwilliams & Davis, 2022).

At the same time, we must be careful not to over-interpret these similarities for several reasons. For one, the TRF is an impulse function that estimates the linear relationship between the acoustic signal and the neural response, thus by definition it is affected by the shape and intensity of the regressor it is trained on. Moreover, TRFs provide an ‘average’ response for the entire regressor they are trained on, and therefore are highly affected by the variability over time across different features of the stimulus. To this point, here we find that the acoustic TRF, that was trained on the entire speech-stimulus, exhibited a double early positive peaks between 0-200ms. However, when separating the analysis according to specific features (e.g., speech rate, opening/closing words), we no longer find a double early peak but rather a single early response peaking at different latencies for different features of the speech. Hence, it is possible that the early double peak in the acoustic model stems from averaging together portions of the speech with different temporal characteristics, and should not be interpreted as a sequence of independent ‘components’. This also highlights the importance of taking this variability into account when modeling the neural response to spontaneous speech. Interestingly, the negative peak in the TRF, seen around 200-400ms, was more stable in latency across the different analyses, suggesting that perhaps it can more reliably be associated with known ERP responses, such as the N400 component, in line with previous speech-tracking studies (Brodbeck et al., 2018, 2022; Broderick et al., 2018, 2019; Donhauser & Baillet, 2020; Gillis et al., 2021). Bearing these caveats in mind, and with the goal of gaining a unified perspective on how findings from more controlled experiments generalize to continuous real-life speech, we now turn to discuss the neural responses to features of spontaneous speech observed here.

### Neural tracking of fillers

The comparison between the TRFs estimated for lexical words vs. fillers indicates a sharp difference in their encoding in the brain. The flat response to fillers was replicated even when parcellating the words according to their length or function. Although fillers clearly serve a communicative function, and some have even evolved from words with lexical meaning (e.g., pragmatic markers that express reduced commitment, such as ‘like’, ‘you know’ (Ziv, 1988)), they do not contribute to the direct comprehension of the narrative and construction of semantic meaning. The substantially flat TRF response observed here for fillers suggests that they are quickly identified as less important, and their encoding is suppressed, perhaps as a means for ‘cleaning-up’ the input from irrelevant insertions before applying syntactic and semantic integrative analysis at the sentence-level.

This finding contributes to an ongoing debate about whether the insertion of fillers into spontaneous speech is a conscious choice of the speaker, just as it is for meaningful words, or if their production is more similar to the insertion of pauses, which are often simply a by-product of production delays (Clark & Fox Tree, 2002; Silber-Varod et al., 2021; Tian et al., 2017). To our knowledge, this is the first study to look directly at the neural response to fillers. Our results suggest that, at least from the listener’s perspective, fillers are treated by the brain more like pauses than words. It is still an open question whether this filtering mechanism is sensitive to the lexicality of the input or rather to the extent to which it contributes to meaning. The fact that no difference was found between function words and content words, which differ in their levels of contribution to meaning, suggests that the lexicality of the input is the important feature. However, since informativeness can be operationalized in various ways which are possibly more accurate than considering only parts of speech, we leave this question open for future research.

### Syntactic boundaries in spontaneous speech

Speech comprehension relies on listeners ability to segment continuous speech into syntactic clauses that serve as basic processing units (Fodor & Bever, 1965; Garrett et al., 1966; Jarvella, 1971). Previous studies have shown that, for highly structured speech stimuli, neural tracking is sensitive to the underlying syntactic structure of the utterances (Ding et al., 2016; Har-shai Yahav & Zion Golumbic, 2021; Kaufeld et al., 2020). However, how this extends to speech that is spontaneous and less well-formed remains unknown. By comparing the neural response to opening vs. closing words of syntactic clauses we sought to find neural markers of detecting the boundaries of syntactic structures ‘on the fly’. Supporting this, here we found that TRFs to closing words had a later latency and larger peak compared to opening words. Importantly, although closing words are longer on average than opening words, this did not seem to explain the observed effects, since no differences were found between TRFs for short vs. long words. Rather, we infer that slightly different neural processing is applied to words that open a new clause vs. the closing word of a clause.

We offer two possible explanations for this result. One explanation pertains to the special status of words at the end of a clause or sentence. As a sentence unfolds, listeners gradually form and update its syntactic representation (J. R. Brennan et al., 2016; Nelson et al., 2017; Pallier et al., 2011). However, only once the sentence ends is it possible to fully integrate across all words to form a complete structure and compute its final syntactic and semantic interpretation. These processes are often referred as ‘wrap up effects’ (Hirotani et al., 2006; Just & Carpenter, 1980; Warren et al., 2009). In the ERP literature, final word in written sentences have been shown to trigger a late positive response, which may be related to the later response seen here for closing words (Friedman et al., 1975; Kutas and Hillyard, 1980). However, since researchers were often advised to avoid analyzing responses to final words (due to wrap up effects), there is very little literature to date directly comparing neural responses to opening vs. closing words, particularly for continuous speech (see Stowe et al., 2018 for a review).

Another possible explanation for the difference in responses to opening vs. closing words relates to the prosodic features of closing words. In speech, syntactic boundaries often align with prosodic breaks (Cooper & Paccia-Cooper, 1980; Klatt, 1975; Strangert, 1992; Strangert & Strangert, 1993) and prosodic cues such as pauses, longer word duration or lower intensity are more prominent on words that close a syntactic unit (Hawthorne & Gerken, 2014; Langus et al., 2012; Vaissière, 1983). Although here opening vs. closing words had similar average intensity and we found that duration alone did not account for the differences, our study was not designed to rule out all prosodic explanations and it is likely that closing words carry other prosodic cues such as pitch deflection and prolonged pauses. Hence, it is possible that the later and larger response seen here for closing words is related to the closure positive shift (CPS), a centro-parietal ERP response observed often at prosodic breaks (Pannekamp et al., 2005; Peter et al., 2014; Steinhauer et al., 1999). Interestingly, the CPS is evoked even without clear prosodic cues, suggesting that it interacts with syntactic boundaries and not simply with acoustic features (Itzhak et al., 2010; Kerkhofs et al., 2007; Steinhauer & Friederici, 2001).

We do not attempt to dissociate the independent contributions of prosodic processing and syntactic processing to online segmentation, and it is highly likely that syntactic and prosodic interpretations are not mutually exclusive, particularly when processing spontaneous speech. Moreover, as noted above, making direct comparisons between ERP components and the TRF responses measured here, is not straightforward. Given the exploratory nature of our study and the lack of extensive literature pertaining to spontaneous speech processing, we offer these results as the basis for data-driven hypothesis, and hope that future research will better characterize the underlying mechanism of the latency shift found here for opening vs. closing words in spontaneous speech.

### The effect of speech rate

When estimating the speech tracking response at the clause-level we found that TRF were significantly earlier for clauses with slow rates vs. those with fast rates. Previous studies have shown that the neural response to speech is highly affected by speech-rate, and that speech tracking and comprehension are poorer for faster speech (Ahissar & Ahissar, 2005; Doelling et al., 2014; Nourski & Brugge, 2011; Verschueren et al., 2022). It is important to note, however, that past research has not studied the natural variability in speech-rate within a given stimulus, but typically artificially manipulate speech rate of the entire stimulus (e.g., by compression).

One interpretation for the earlier response found here for slower speech could be acoustic, since the envelope of slower speech can also have a shallower-slopes at acoustic edges (Oganian et al., 2022). These have been shown in a previous study to affect the amplitude of the TRF response, which was larger for steeper slopes (Oganian & Chang, 2019). Another recent study found that faster speech induced larger TRF amplitude as well as an increased TRF latency, interpreting this finding as stemming from the more time that is needed to process fast speech (Verschueren et al., 2022). However, another interpretation may relate not to durational features of words, which could be longer in slow speech, but to the frequent disfluencies that are found in slow speech. Pauses and fillers allow more time for the listener to adjust their expectations and predict upcoming words in the utterance (Clark & Fox Tree, 2002; Fox Tree, 2001; Fraundorf & Watson, 2011; Goldman-Eisler, 1958a, 1958b; Watanabe et al., 2008). Hence, the earlier latency of the TRF may reflect the effects of predictive processing that may be easier to apply to slower speech (Zion Golumbic et al., 2012).

### Summary

As speech-processing research moves towards understanding how the brain encodes speech under increasingly realistic conditions, it is important to recognize that the type of speech we hear daily is linguistically imperfect. The current study offers insight into some of the ways that these imperfections are dealt with in the neural response and demonstrates the robustness of the speech processing system for understanding spontaneous speech despite its disfluent and highly complex nature. The mechanistic and linguistic insights from this proof-of-concept study provide a basis for forming specific hypotheses for the continued investigation of how the brain deals with the natural imperfections of spontaneous speech.

