## supplemental data for "“Um…, it’s really difficult to… um… speak fluently”: Neural tracking of spontaneous speech"

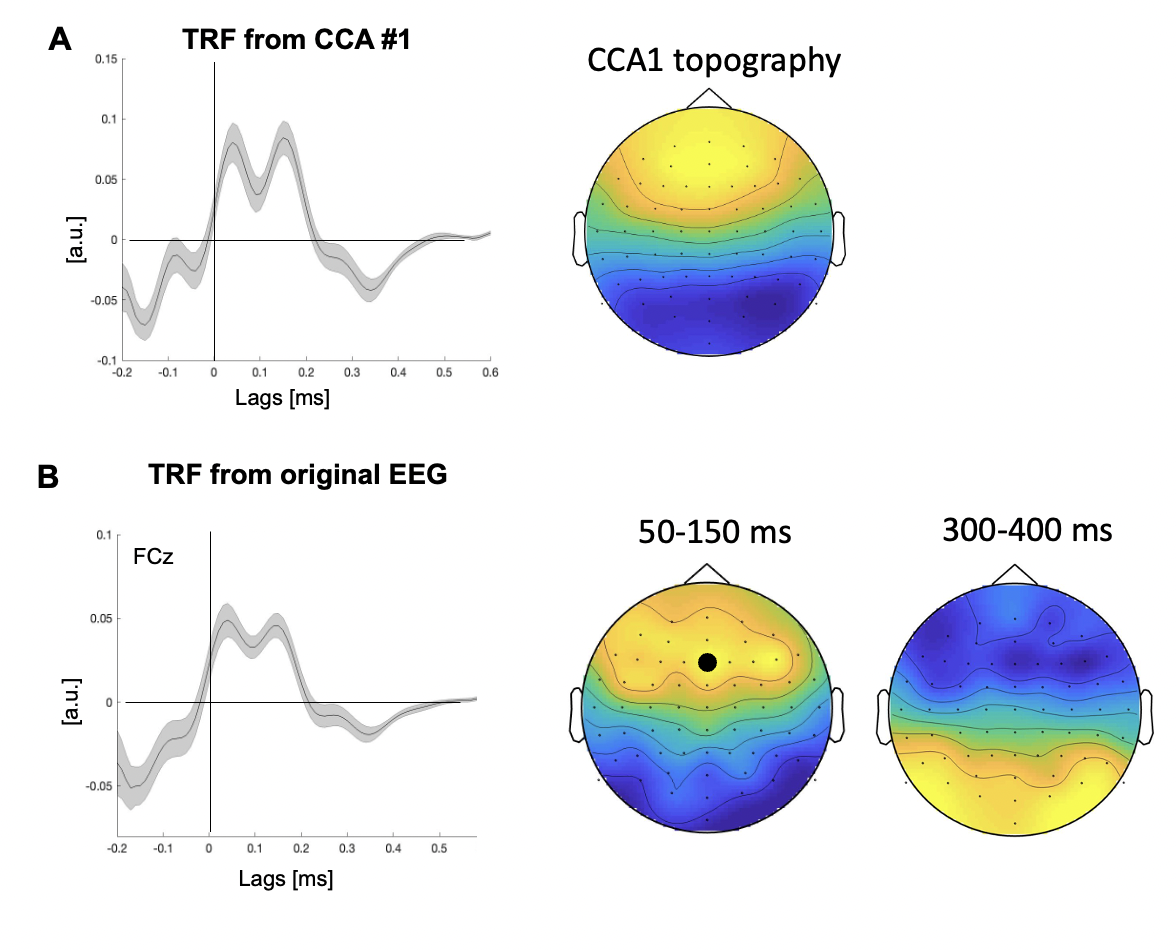
 **Figure S1.** Comparison of the TRF estimated for **(A)** the first CCA component (CCA #1) vs. **(B)** TRFs estimated using the original EEG data (64-channels, bandpass filtered 1-20Hz, shown here for electrode FCz). Both TRFs share a common time-course and the scalp topography of CCA #1 matches the scalp topography of the two peaks in the original EEG-TRF (early positive peak between 50-150ms and late negative peak between 300-400ms). This indicates that CCA #1 reliably captures the portion of the neural signal that tracks the speech and that this method can be used for dimension-reduction without substantial data loss.

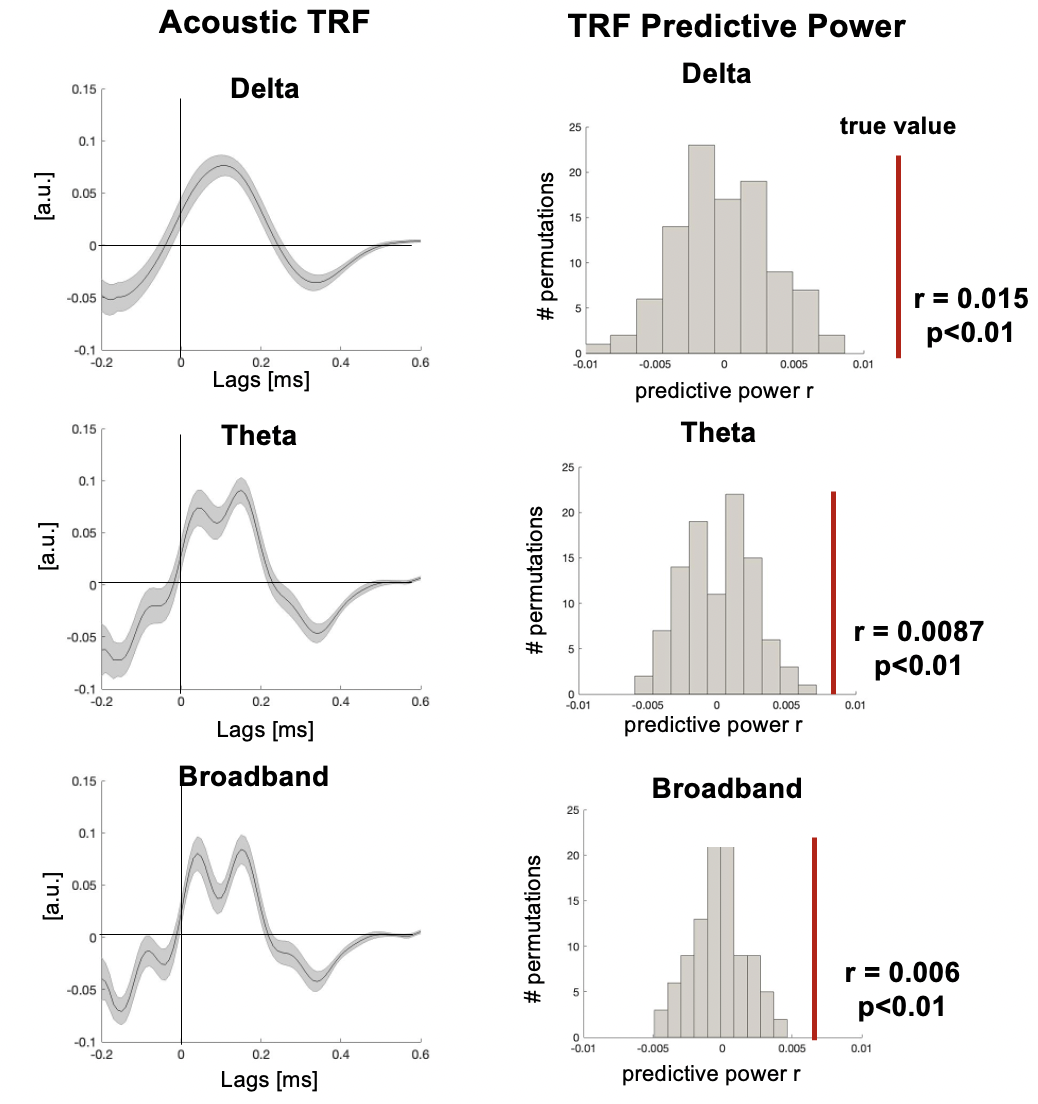
 **Figure S2. Left:** The TRF estimated for CCA component #1 for different frequency bands (delta: 1-3Hz, theta 4-8Hz, broadband 1-20Hz). **Right:** The predictive power of the TRFs shown on the left (red line), relative to the null-distribution of predictive power from 100 randomly shuffled S-R combinations. In all cases, the predictive power is highly significant, however the predictive-power values themselves (for the null-distribution and the real data) are affected by filtering.
